# Sex-dependent effect of inflammatory pain on negative affective states is prevented by kappa opioid receptors blockade in the nucleus accumbens shell

**DOI:** 10.1101/2023.04.22.537926

**Authors:** J.D. Lorente, J. Cuitavi, L. Rullo, S. Candeletti, P. Romualdi, L Hipólito

**Author notes:** Conceptualisation: Jesús D. Lorente and Lucía Hipólito. Methodology: Jesús D. Lorente, Javier Cuitavi, Laura Rullo, Sanzio Candeletti, Patrizia Romualdi and Lucía Hipólito. Formal analysis: Jesús D. Lorente, Laura Rullo, Sanzio Candeletti, Patrizia Romualdi and Lucía Hipólito. Investigation: Jesús D. Lorente, Javier Cuitavi, Laura Rullo. Writing: Jesús D. Lorente, Javier Cuitavi, Laura Rullo, Sanzio Candeletti, Patrizia Romualdi and Lucía Hipólito. Resources: Sanzio Candeletti, Patrizia Romualdi and Lucía Hipólito. Supervision: Sanzio Candeletti, Patrizia Romualdi and Lucía Hipólito All authors contributed to the article and approved the submitted version. **Conflict of interest and ethical considerations:** The authors declare that the research was conducted in the absence of any commercial or financial relationships that could be construed as a potential conflict of interest. The animal study was reviewed and approved by the Ethics Committee in Experimentation and Animal Welfare of the University of Valencia and the local Government of Valencia (Conselleria d’Agricultura, Desenvolupament Rural, Emergéncia Climática i Transició Ecológica). **Data availability statement** The raw data supporting the conclusions of this article will be made available by the authors, without undue reservation.

## Abstract

**Background and purpose:** Psychological disorders, such as anxiety and anhedonia are pain comorbidities, however how pain affects male and female individuals and through which mechanism is not well understood. Previous research show pain-induced alterations in the dynorphinergic pathway in the mesocorticolimbic system (MCLS) together with a relationship between corticotropin-releasing system and dynorphin release in the MCLS. Here, we analyse the sex and time course-dependent effects of pain on negative affect. Additionally, we study the implication of dynorphinergic and corticotropin releasing factor involvement in these pain related behaviours.

**Experimental approach:** We used behavioural pharmacology and biochemical tools to characterize negative affective states induced by inflammatory pain in male and female rats, and the alterations in dynorphinergic and corticotropin systems in the MCLS.

**Key results:** Female rats showed a persistent anxiety-like together with a reversible anhedonia-like behaviours derived from inflammatory pain. Additionally, we found alterations of in both dynorphin and corticotropin releasing factor in NAc and amygdala that suggest sex-dependent dynamic adaptations. Finally blockade on the kappa opioid receptor in the NAc confirmed its role in pain-induced anxiety-like behaviour in female rats.

**Conclusions and implications:** Our results show sex and time dependent anxiety- and anhedonia-like behaviours induced by the presence of pain in female rats. Furthermore, we replicated previous data pointing to the KOR/dyn recruitment in the NAc as key neurological substrate mediating these behaviours. This research encourages the study the mechanisms underlying these behaviours, to better understand the emotional dimension of pain.

## 1. Introduction

Pain is suffered by 30% of adults in developed countries (Sá et al., 2019). The therapeutical plan of chronic pain has traditionally been focused on alleviating the symptomatology. In fact, psychological comorbidities have been overlooked and, in the vast majority, no treatment has been applied. During the last years, pain management has become a multidisciplinary area including different specialist. By applying different psychological therapies in combination with pharmacotherapy and other pain-relief techniques, patients have improve their quality of life, easing their pain sensation, diminishing the possibility of disability, and reducing fear-avoidance behaviours (Petrucci et al., 2022). Interestingly, men and women show different pain physiopathology resulting in discrepancies in pain feeling, response to pain killers and psychological comorbidities (Edwards et al., 2003; Pieh et al., 2012; Popescu et al., 2010; Riley et al., 1998). Nevertheless, since females have traditionally been excluded in research (Burek et al., 2022), the lack of sex-based studies could have led to undertreat pain-related conditions in women, promoting chronic pain and the development of neuropsychiatric comorbidities (Cáceda et al., 2021; Jakubczyk et al., 2016). Moreover, a metanalysis performed by Burek and collaborators showed that the classical behavioural tests of anxiety- and depression-like behaviours in the presence of pain lead to contradictory results depending on the strain, source, sex of the animal and testing time after the induction of pain (Burek et al., 2022)

Recent reports highlighted that pain induces alterations in the mesocorticolimbic system (MCLS) in both, kappa opioid receptor (KOR)/dynorphin (DYN) and corticortropin-releasing factor (CRF) systems (Fu & Neugebauer, 2008; Ji et al., 2007; Markovic et al., 2021; Massaly et al., 2019; Mazzitelli et al., 2022; Navratilova et al., 2019). Along with other effects within the MCLS, the KOR/DYN system recruitment in the nucleus accumbens (NAc) has shown to mediate negative affective states induced by inflammatory and neuropathic pain (Liu et al., 2019; Massaly et al., 2019). Additionally, presence of inflammatory pain has led to alterations in the CRF 1 receptors (CRF1R) transmission in central amygdala (CeA) (Fu & Neugebauer, 2008; Ji et al., 2007). In fact, CRF1R activation in CeA is necessary to develop pain-related anxiety-like behaviour, and the activation of KORs in CeA disinhibits CRF1R producing pain-related anxiety-like behaviour (Hein et al., 2021; Ji et al., 2007; Mazzitelli et al., 2022). Furthermore, the optoinhibition of CRF containing neurons in CeA is sufficient to impair anxiety-like behaviour induced by pain (Hein et al., 2021; Mazzitelli et al., 2022). Interestingly, KOR blockade in CeA controls pain-related behaviour in a model of functional pain by restoring synaptic inhibition of CeA-CRF neurons, demonstrating that KOR and CRF systems are closely related in pain and its comorbidities (Yakhnitsa et al., 2022). However, most of these cited papers have obtained these results only in male rodents leaving females understudied.

In order to shed light on the time course and sex-dependent effects of pain-induced comorbidities, here we will use male and female rats to be tested in classical anxiety/depression behavioural assays for a total period of three weeks. Additionally, we will study the CRF and KOR/DYN systems alterations in the MCLS that may be implicated in the observed behaviours.

## 2. Methodology

### 2.1. Animals and pain model

A total of 124 Sprague-Dawley rats (Envigo, Barcelona, Spain) of 7-8 weeks old at the beginning of the behavioural experiments, 100 females (170-200 g) and 24 males (260-300 g) were used. Animals were housed individually in standard plastic cages (42 x 27 x 18 cm^3^) provided with shredded aspen bedding (Teklad, Barcelona, Spain) and cotton enrichment (iso-BLOXTM; Teklad) and were maintained in 12/12 hours light/dark cycles (light off at 07:00 am, light on at 07:00 pm), at 23 ± 1 ° C and 60% humidity. All behavioural tests were conducted during the dark cycle and at least 2 hours after the lights turn off. Food and tap water were available *ad libitum* throughout the experimental period. The protocols used were approved by the Animal Care Committee from the University of Valencia and authorised by the regional government, and the studies were performed in strict accordance with Spanish laws (RD 53/2013) and European Directives (EC 2010/63). Animals were sacrificed by isoflurane overdose for further biochemistry analysis.

Rats received a subcutaneous injection of 0.1 mL of the complete Freund adjuvant (CFA; diluted at 1:2 in sterile saline to create an emulsion) in the hindpaw (Hipolito et al., 2015). This animal model of pain based on the CFA administration has been broadly used for replicating aspects of arthritis (Fischer et al., 2017).

### 2.2. Experimental design

#### 2.2.1. Experiment 1: inflammatory pain effect on negative affective states, corticortropin releasing factor system and Kappa/dynorphin system in the Nucleus Accumbens and amygdala

##### 2.2.1.1. Behavioural experiments design and brain collection

To study the time course and the gender effect of negative affective states induced by inflammatory pain we have selected a combination of different behavioural approaches. One week prior to CFA administration, we obtained basal nociceptive measures by means of Von Frey test (VFT) together with basal measures of the sucrose preference test (SPT). During the following 18 days after complete Freund Adjuvant (CFA) administration, animals were repetitively tested in the VFT, light-dark box test (LDB) and SPT (n=12-14/group). Finally, 18 days after the CFA administration, rats were sacrificed to collect their brains. Details of this protocol and timeline are summarized in figure 1A and figure S1.

**Figure 1.**
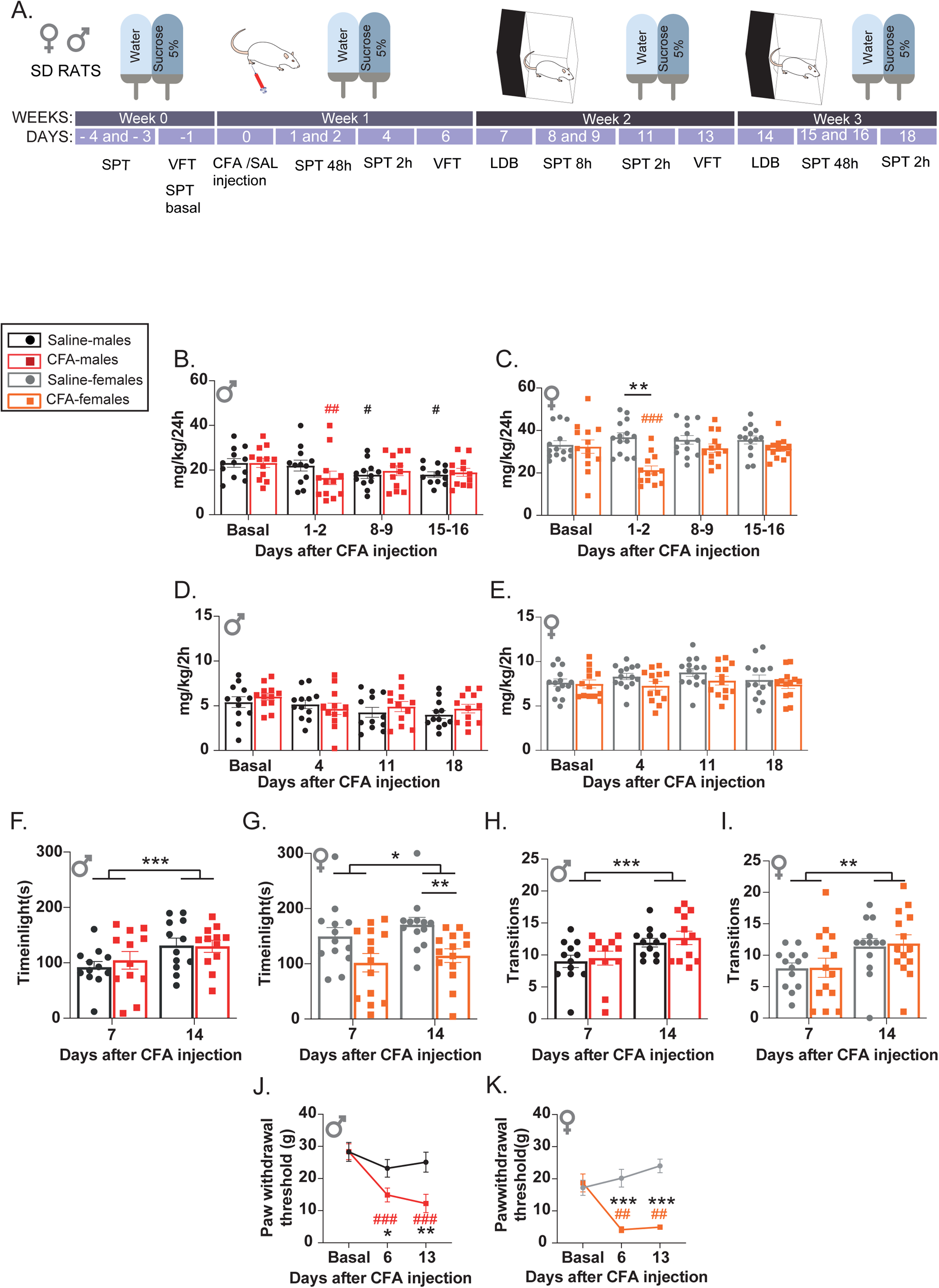
Anhedonia- and anxiety-like behaviour induced by inflammatory pain in male and female rats. A) Timeline of the experimental design B) Sucrose consumption during 48 hours in male, expressed as mean ± SEM of mg/kg/24 hours C) Sucrose consumption during 48 hours in female, expressed as mean ± SEM of mg/kg/24hours D) Sucrose consumption during 2 hours in male, expressed as mean ± SEM of mg/kg/2 hours E) Sucrose consumption during 2 hours in female, expressed as mean ± SEM of mg/kg/2 hours F) Time (seconds) in light chamber in the light dark box test (LDB) in male, expressed as mean ± SEM G) Time (seconds) in light chamber in the LDB in male, expressed as mean ± SEM H) Number of transitions between chambers in the LDB in male, expressed as mean ± SEM I) Number of transitions between chambers in the LDB in female, expressed as mean ± SEM J) Mechanical sensitivity threshold in male, expressed as mean ± SEM of paw withdrawal in grams K) Mechanical sensitivity threshold in female, expressed as mean ± SEM of paw withdrawal in grams. Black and red bars and symbols represent saline- and CFA-treated male respectively, and grey and orange bars and symbols represent saline- and CFA-treated female respectively. *, ** and *** denotes significant differences between groups (two-way ANOVA for repeated measures followed by Bonferroni multiple comparisons test, p< 0.05, p< 0.01 and p< 0.001, respectively), and #, ## and ### denotes significant differences within groups compared with the basal measures (two-way ANOVA for repeated measures followed by Bonferroni multiple comparisons test, p< 0.05, p< 0.01 and p< 0.001, respectively). Abbreviations: SPT, sucrose preference test; VFT, Von Frey test; CFA, Complete Freund’s Adjuvant; SAL, saline; LDB, light-dark box test.

##### 2.2.1.2. Biochemicals analysis

###### Blood and brain samples collection and preparation

Immediately after the overdose of isoflurane, blood samples, were obtained by heart puncture and placed in plastic tubes, which were flash frozen in contact with dry ice. Immediately, after collecting blood samples, we extracted brains which were also flash frozen in dry ice.

The plasma collection was performed the same day the ELISA assays were performed. To extract plasma, we first defrosted the blood samples in contact with ice. Once all the samples were thawed, we kept them 30 min in the ice. Next, we centrifugated them for 10 minutes at 33500 G, and we collected the supernatant.

Tissue from amygdala and NAc was collected by punching in brain slices of 61.2 and 61.6 µm respectively. The bregma coordinates of the brain slices were from 2.52 to 0.96 mm for NAc, and from -2.04 to -3.24 mm for amygdala (Figure S2).

###### RNA isolation and qRT-PCR

Total RNA was isolated using TRIZOL reagent (Life Technologies, USA) according to the method of Chomczynski and Sacchi (1987). The integrity of the samples was verified by 1% agarose gel electrophoresis and the RNA amount in each sample was measured by spectrophotometry, as described (Caputi, et al., 2021a).

RNA samples were subsequently subjected to DNase treatment and converted to cDNA using the GeneAmp RNA PCR kit (Life Technologies, Italy) according to the manufacturer’s protocol.

The qRT-PCR analysis was performed on a StepOne Real-Time PCR System (Life Technologies, Monza, Italy) using SYBR Green PCR MasterMix (Life Technologies, Italy), as previously reported (Caputi, et al., 2021b). Relative expression of the different gene transcripts was calculated by the Delta-Delta Ct (DDCt) method and converted to relative expression ratio (2-DDCt) for statistical analysis (Livak & Schmittgen, 2001). All data were normalised to the housekeeping gene glyceraldehyde-3-phosphate dehydrogenase (GAPDH). The primers, designed using Primer 3 (Rozen & Skaletsky, 2000) were used for PCR amplification.

pDYN forward 5′− CCTGTCCTTGTGTTCCCTGT-3′; pDYN reverse 5′-AGAGGCAGTCAGGGTGAGAA-3′; KOR forward 5′-TTGGCTACTGGCATCATCTG-3′; KOR reverse 5′-ACACTCTTCAAGCGCAGGAT-3′; CRF forward 5′-GCAGCGGGACTTCTGTTGA-3′; CRF reverse 5′-CGCAGCCGTTGAATTTCTTG-3′; CRFR1 forward 5′-TGCCAGGAGATTCTCAACGAA-3′; CRFR1 reverse 5′-AAAGCCGAGATGAGGTTCCAG-3′; GAPDH forward 5′-AGACAGCCGCATCTTCTTGT-3′; GAPDH reverse 5′-CTTGCCGTGGGTAGAGTCAT-3′.

###### Protein extraction

For protein analysis, we used a previous published protocol of protein extraction (Lorente et al., 2022). 0.5 mL of cold lysis buffer (1% IGEPAL CA-630, 20mMTris-HCl pH 8, 130MNaCl, 10mMNaF, and 1% proteases inhibitor cocktail [Sigma, Darmstad, Germany]) was used to homogenate 250 mg of brain tissue, thus the volume of the lysis buffer was adapted to the real quantity of tissue in each sample. Then we maintained the samples in ice for 30 minutes, before centrifugate at 13000 G for 15 minutes at 4 ° C and collect the supernatant. Furthermore, we determined the concentration of the lysates by using a Bradford protein assay kit (Bio-Rad).

###### ELISAs

Two different ELISA kits (Cusabio, Houston, USA) were used to detect the levels of DYN in amygdala and NAc samples (#REF CSB-E13294r) and cortisol in plasma (#REF CSB-E05112r), according to kit protocols.

###### Western blot

We used a protocol previously used in our laboratory (Cuitavi et al., 2021). 1-mm acrylamide gels at 10% with 15 wells were used for the electrophoresis. We mixed 20 µg of total protein, loading buffer (350 mM Tris pH 6.8, 30% glycerol, 30% mercaptoethanol, 100g/L sodium dodecyl sulphate, and 200 mg/L bromophenol blue) and water to obtain the same total protein and volume per sample, and they were heated at 70°C for 20 minutes. We used the Bio-Rad Mini-PROTEAN buffer system (6 g/L Trizma base, 2.88 g/L glycine, and 20 g/L sodium dodecyl sulphate) to perform the electrophoresis at 120 V for 80 to 90 minutes. Once, the proteins were separated by sodium dodecyl sulphatepolyacrilamide gel electrophoresis (SDS-PAGE), we transferred it to nitrocellulose membranes (Bio-Rad) with the appropriate buffer (3 g/L Trizma base, 1.44 g/L glycine, and 20% methanol) and a semidry system (Bio-Rad Trans-Blot TurboTM) for 25 minutes at 25 V buffer.

Then, membranes were blocked using 5% non-fat dried milk in TBS-Tween 20 (TBS-T) 0.1% (20 mM Tris and 500 mM NaCl pH 7.5) for 1 hour, and just before they were incubated with the corresponding primary antibody (rabbit anti-KOR 1:2000, REF #44-302G, Thermo Fisher; goat anti-CRFR1 1:1800, REF #PA5-18801, Thermo Fisher; rabbit anti-GAPDH 1:2000, Thermo Fisher) overnight at 4° C. After that, membranes were washed 3 times with TBS-T 0.1% and were incubated for an hour at room temperature with the secondary antibody (Goat anti-Rabbit IgG (H+L) HRP conjugated at 1:2000 for GAPDH and KOR (REF #31460, Thermo Fisher; Donkey anti-Goat IgG (H+L) HRP conjugated 1:2000 for CRF1R, REF #A15999, Thermo Fisher). Finally, we developed the membranes with chemiluminescence using CheLuminate-Horseradish peroxidase (HRP) PicoDetect (Panreac, Barcelona, Spain), and images were captured with the ChemiDoc XRS1 System (Bio-Rad) and quantified using ImageJ software. The grey intensity of the bands was normalised dividing the values of the protein of interest by the GAPDH values. The relative protein levels were expressed as mean ± SEM.

#### 2.2.2. Experiment 2: effect of norbinaltorphimine administration in the nucleus accumbens medial shell on complete Freund adjuvant–induced negative affective states in female rats

##### 2.2.2.1. Behavioural experiment protocol

To study the influence of KOR/DYN system in pain-induced negative affective states in NAc of female rats, the same protocol of experiment 1 was used except for the additional bilateral infusion of the kappa opioid receptor antagonist norbinaltorphimine (NorBNI) into the NAc medial shell. The week prior to the CFA administration after the basal measures for VFT and SPT were obtained, animals were injected with NorBNI or artificial cerebral spinal fluid (aCSF) into the NAc medial shell. The following 14 days after CFA administration animals were tested following the VFT, LDB and SPT paradigms. The timeline followed is summarised in figure 4A and figure S3A.

##### 2.2.2.2. Surgery and norbinaltorphimine intra-accumbal administration

All surgeries were performed under isoflurane anaesthesia (1.5-2 minimum alveolar concentration, MAC) and under aseptic conditions. Before anesthetising the animals for the surgeries, they received injections (s.c.) of 1.8 mg/kg enrofloxacin (Bayer) and 2.5 mg/kg of carprofen (Pfizer). Then, we anesthetised animals in a induction chamber and used 0.1% topical lidocaine in the surgical area and in the ears before fixing animals to the stereotaxic frame (Stoelting, Wood Dale, USA). The skull of the rats were exposed, and a craniotomy was bilaterally performed above the posteromedial NAc shell. By means of a stainless steel microinjector (33-gauge) attached to a PE-10 tubing and a 25-mL Hamilton syringe mounted on a syringe pump (Kd Scientific, Holliston, USA) we injected aCSF (0.5 uL per side) or NorBNI (2 ug per hemisphere in 0.5 uL, REF #N1771, SigmaAldrich, St. Louis, USA) in the following coordinates: +0.96 mm anteroposterior, ±0.8 mm mediolateral, and -6.2 mm dorsoventral from bregma in a flat skull position (Massaly et al., 2019). Finally, we covered the craniotomies with bone wax (Ethicon, Cincinnati, USA) and the skin of each animal was sutured with a nylon monofilament suture (Ethicon, Cincinnati, USA). Rats were allowed to recover in a box provided with a heat pad and were closely monitored until they fully recovered from the anaesthesia. During the days following the surgery, rats were daily examined and received injections (s.c.) of 1.8 mg/kg enrofloxacin and 2.5 mg/kg of carprofen once a day the following 2 days.

##### 2.2.2.3. Microinjection placement assessment

At the end of the protocol, rats were sacrificed under isoflurane overdose and brains were removed and rapidly frozen in dry ice; 40 µm-thick coronal slices of the NAc shell were obtained using a cryostat (Micron, Boise, USA) and stained with a cresyl violet (Sigma, St. Louis, USA) protocol to verify proper probe placement as we have used previously (Campos-Jurado et al., 2020). Only animals that received a bilateral injection into the NAc medial shell were included in the analysis (figure S3B).

### 2.3. Behavioural procedures

#### Sucrose preference test

For SPT, a slightly modified protocol of a reported procedure (Schalla et al., 2020) has been used. Briefly, animals had free access to 5% sucrose diluted in tap water along with tap water twice a week, in two different types of sessions. The first session lasted 48 hours (divided in 2 consecutive sessions of 24h, since bottles needed to be weighted and refilled), and was followed by a period of 24h without access to sucrose. Following, the second session started and lasted for 2 hours. After obtaining a basal measure the week prior to the pain induction, we repeated the protocol every week until the end of the experiment. All measurements were expressed as the mean ± SEM in mg of sucrose/rat weight in kg/24h or mg/kg/2h.

#### Light-dark box test

The LDB apparatus consists of a box (64 x 48 x 24 cm^3^) that is divided in 2 different chambers: light chamber (two-thirds of the total size) and dark chamber (one-third of the total size). The light and dark chambers are connected by a squared opening (8 x 8 cm^2^). The light chamber, which is uncovered, has white walls, and is illuminated by white light (400-lux). The dark chamber has black walls and has no appreciable illumination at the centre of the chamber (≤ 2 lux). After at least 5 min of habituation, animals were gently placed at the centre of the light box, with the head facing opposite to the squared opening, and then animals explored freely for 5 minutes. Spontaneous behaviours of the rats were videotaped in the light box during the test for its posterior analysis. The test was performed during the dark cycle, at least 2 hours after the lights went off. The analysed parameters were the time spend in the light box (seconds, s) and the number of transitions between chambers. All the measurements were expressed as the mean ± SEM.

#### Von Frey test

The VFT was used to assess the mechanical nociceptive thresholds before (basal) and after the CFA injection. After 5-10 min of habituation to the apparatus, Von Frey filaments (Aesthesio®) were manually applied to the injected hind paw with the simplified up-down method (Bonin et al., 2014). Results were expressed as the mean ± SEM of mechanical sensitivity threshold (in grams, g).

### 2.4. Statistical analysis

After testing the normality of distribution data (Shapiro-Wilk test) and homogeneity of variance, unpaired Student’s t test, Welch’s t test, Mann-Whitney U test were used to analyse gene expression, ELISAs and western blot results. Statistical analysis was performed using GraphPad Prism 9 software (San Diego, USA) or IBM SPSS statistics v24 software. For the behavioural data, we used the ANOVA for repeated measures followed by Bonferroni multiple comparisons as *post hoc* test when appropriate. Results are expressed as mean ± SEM and the statistical significance was set at p < 0.05. All the p values of the statistic tests are reported in Table 1 and the p values from the *post hoc* analysis are described in the figure legends.

## 3. Results

### 3.1. Sex- and time-dependent anhedonia and anxiety-like behaviours induced by inflammatory pain

Pain highly impacts patients’ quality of life by promoting negative affective states (Pieh et al., 2012; Popescu et al., 2010). Thereby, in this study we assess the time course of the development of these negative affective states in an inflammatory pain model based on CFA injection. To this end, we have monitored anhedonia and anxiety-like behaviours (SPT and LDB respectively) for several weeks (fig. 1A and S1).

We selected the SPT as a behavioural approach to measure anhedonia. First, we analysed the total sucrose consumption (mg/kg/24h) in the 48h test on post-CFA injection days 1-2, 8-9 and 15-16, followed by the analysis of the total sucrose consumption in the 2h test on post-CFA injection days 4,11 and 18 (fig 1A). In male rats, the statistical analysis of the 48h test revealed significant time (F (3, 66) = 6.636, p = 0.001, partial η^2^= 0.966) and the pain x time interaction effect (F (3, 66) = 3.709, p = 0.016, partial η^2^= 0.783; table 1) but no effect for the pain variable (F (1, 22) = 0.69, p=0.795, partial η^2^= 0.057). Thus, the development of the inflammatory condition decreased the sucrose consumption along the experimental procedure for both pain and no pain group. Their sucrose intake at days 1-2 for pain group, and at days 8-9 and 15-16 for control group compared with their basal measures in the 48h test was significantly lower (fig 1B, Bonferroni multiple comparisons intra-subjects: p= 0.04; p= 0.17 and p= 0.034, respectively). The analysis of the 2h access to sucrose sessions by the repeated measures ANOVA only detected time dependent effect (F (3, 66) = 8.886; p < 0.001, partial η^2^= 0.993), but no differences between groups were found in the Bonferroni multiple comparisons test (fig. 1D). However, female rats have shown a different pain-induced anhedonia-like behaviour profile (fig. 1C). Indeed, the ANOVA detected pain (F (1, 25) = 7.688, p=0.01, partial η^2^= 0.760), time (F (3, 75) = 3.496, p=0.02, partial η^2^= 0.759) and the interaction pain x time (F (3, 75) = 7.819, p= 0.001, partial η^2^= 0.986) significant effects. In fact, the post-hoc analyses showed that CFA-females reduced their sucrose consumption during days 1-2 compared with saline-females (p= 0.0001) and with their own basal intake (p= 0.003). Interestingly, CFA-females recovered from the anhedonia showed in the 1-2 days post-CFA injection; indeed, no changes in total sucrose intake were observed in the next two sessions in comparison with the respective controls (comparisons with basal: p_8-9_= 1.000 and p_15-16_= 1.000; comparisons with 1-2 days after CFA-injection: p_8-9_= 0.001 and p_15-16_= 0.0001). The 2 h tests showed no alterations in the sucrose intake derived from the presence of pain during the experimental procedure since no statistical differences were observed in none of the studied variables (fig.1E, table 1).

Second, anxiety-like behaviours induced by inflammatory pain were studied with the LDB test by analysing the time spent in the light chamber (Figures 1F and 1G). The ANOVA for repeated measures detected only an effect of time in both measures, time in light (F (1, 22) = 19.284; p= 0.0001, partial η^2^= 0.987) and number of transitions (F (1, 22) = 19.284; p= 0.0001, partial η^2^= 0.987) in the case of males (Fig 1F and 1H), meaning that inflammatory pain has no effect on anxiety-like behaviour. Indeed, males showed an increase of the time in the light box and the number of transitions in the second week when compared to the first week probably due to habituation.

Interestingly, the analysis of data from female rats, revealed an effect for pain (F (1, 25) = 7.287; p = 0.012) and time (F (1, 25) = 4.355; p = 0.047, η^2^= 0.519) but not for the interaction pain x time (F (1, 25) = 0.278; p = 0.603, η^2^= 0.080; fig 1G). Thus, the post-hoc analysis revealed that CFA-treated females reduced their time spent in the light chamber during the second week compared with non-pain group (p= 0.05). The analysis of the number of transitions in the LDB, showed no differences in pain, time or the interaction of these two variables for male and female rats, indicating no differences in the general motor activity and exploration ability of the CFA-treated rats.

Finally, the mechanical sensitivity threshold was assessed along the experimental procedure by using the Von Frey test. Figures 1J and 1K show that CFA-treated male and female rats maintained a low threshold after the CFA injection. The statistical analysis showed an effect for pain, time and pain x time interaction in both sexes (Male rats: pain F(1,22)= 6.223, p= 0.665, η^2^=; time F(2,44)= 4.148, p= 0.022, η^2^= 0.932; pain x time F(2,44)= 11.538, p= 0.0001, η^2^= 0.639; female rats: pain F(1,25)= 56.140, p=0.0001, η^2^= 1; time F(2,50)= 3.252, p=0.047, η^2^= 0.265; pain x time F(2,50)= 11.783, p= 0.0001, η^2^= 0.975). Post-hoc analyses revealed that both male and female rats in pain showed significant lower threshold at day 6 and 13 when compared with their own basal and with no pain group (Male rats: p_6_< 0.027 and p_13_< 0.006; Female rats: p_6_< 0.001 and p_13_< 0.001)

### 3.2. Inflammatory pain influence on corticotropin releasing factor (CRF), CRF receptor 1 (CRFR1) and blood cortisol levels

CRF signalling is increased as a result of the presence of stress activating the hypothalamus-pituitary-adrenal axis (HPA) to produce a neurohormonal response. Chronic stress and negative affect derived from this stress affects the HPA and the functioning of some brain areas such as amygdala or nucleus accumbens. Activation of the HPA system leads to increased corticosteroids blood levels to respond to a threat or stressful event, but finally this increase exerts negative feedback to the HPA system to limit this response (Herman et al., 2016; Koob et al., 2014). Thereby, we measured mRNA and protein levels of CRF and CRFR1 in the NAc and the Amy by qRT-PCR and western blot. In addition, we also measured blood cortisol levels by ELISA (fig. 2).

**Figure 2.**
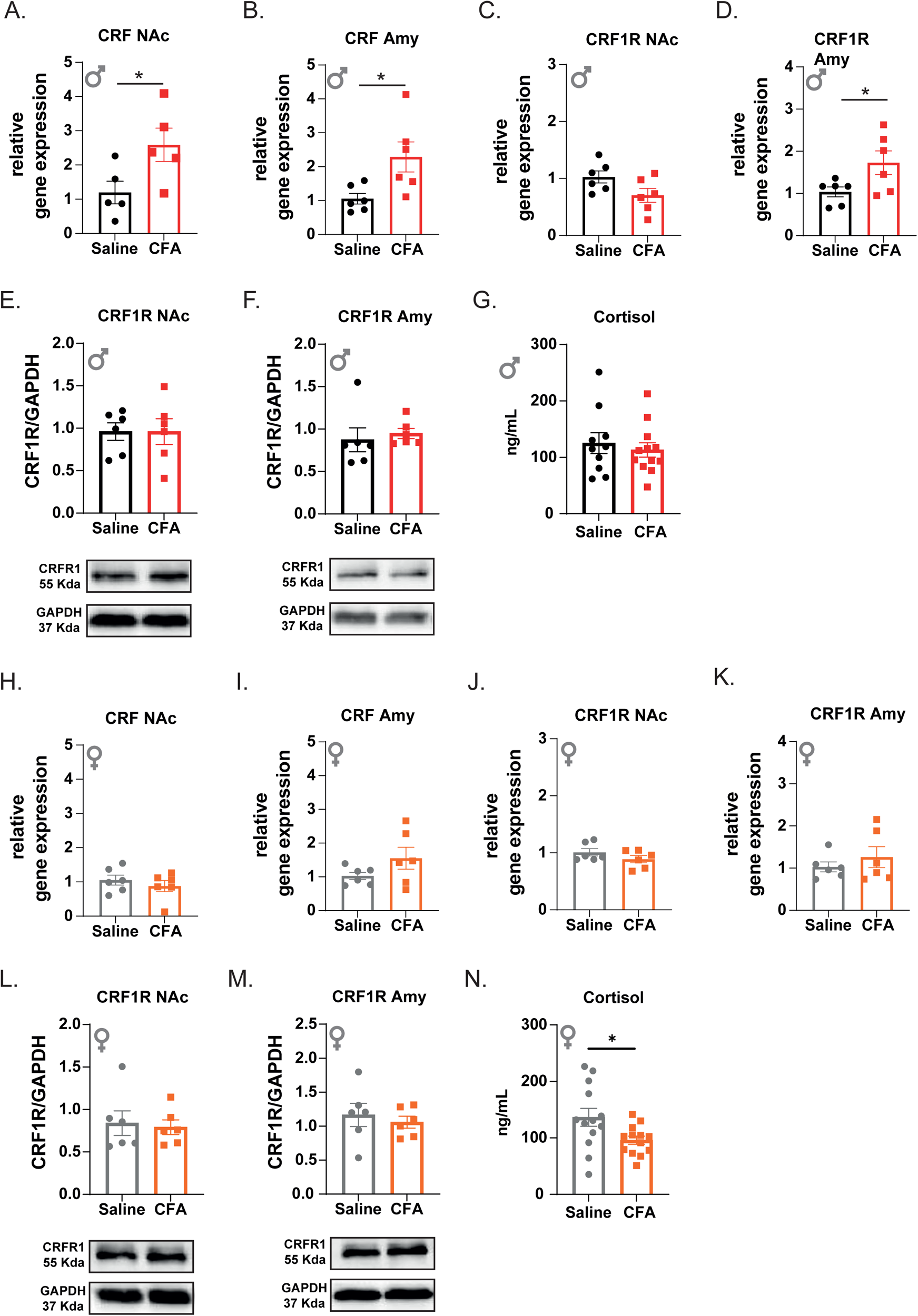
Corticotropin-releasing factor system alterations induced by inflammatory pain. A) Mean ± SEM of CRF mRNA relative gene expression in NAc of male rats. B) Mean ± SEM of CRF mRNA relative gene expression in Amy of male rats. C) Mean ± SEM of CRF1R mRNA relative gene expression in NAc of male rats. D) Mean ± SEM of CRF1R mRNA relative gene expression in Amy of male rats. E) Mean ± SEM of CRF1R protein expression in NAc of male rats (CRF1R/GAPDH arbitrary units). F) Mean ± SEM of CRF1R protein expression in Amy of male rats (CRF1R/GAPDH arbitrary units). G) Plasma cortisol levels in male rats, expressed as mean ± SEM of ng/mL. H) Mean ± SEM of CRF mRNA relative gene expression in NAc of female rats I) Mean ± SEM of CRF mRNA relative gene expression in Amy of male rats. C) Mean ± SEM of CRF1R mRNA relative gene expression in NAc of female rats. J) CRF1R expression in NAc in female, expressing as mean ± SEM of relative gene expression. K) Mean ± SEM of CRF1R mRNA relative gene expression in Amy of female rats. L) Mean ± SEM of CRF1R protein expression in NAc of female rats (CRF1R/GAPDH arbitrary units). M) Mean ± SEM of CRF1R protein expression in Amy of female rats (CRF1R/GAPDH arbitrary units). O) Plasma cortisol levels in female rats, expressed as mean ± SEM of ng/mL. Black and red bars represent saline- and CFA-treated male respectively, and grey and orange bars represent saline- and CFA-treated female respectively. * Denotes significant differences between groups (t-student for independent samples or Welch’s t test p < 0.05).

mRNA levels of CRF in NAc and Amy where significantly increased in CFA-treated male rats (fig 2A, p= 0.045; 2B, p=0.038) together with an increase in the mRNA of CRFR1 only in Amy and not in NAc (fig 2C, p= 0.075; fig 2D, p= 0.046) at the endpoint of the experimental protocol. However, further analysis of the CRFR1 protein expression showed that this altered expression in the mRNA had no effect on the local receptor expression (Figure 2E, p= 0.999 and figure 2F, p= 0.093). Finally, blood cortisol levels, at this time point, were not altered in CFA-treated male rats (fig 1G, p= 0.589).

Very interestingly, we did not observe any changes in the levels of the CRF or CRFR1 mRNAs or proteins in the NAc or in the Amy of CFA-female rats (figure 2H, p= 0.445; figure 2I, p= 0.177; figure 2J, p= 0.394; figure 2K, p= 0.42; figure 2L, p= 0.818; figure 2M, p= 0.590). However, as shown in figure 2N, females with inflammatory pain presented a significant lower blood cortisol level than saline-treated females (p = 0.033).

### 3.3. Dynorphin/kappa opioid receptors mRNA and protein levels alterations in the mesocorticolimbic system induced by inflammatory pain

The dynorphinergic system, which comprises the KOR and their ligand DYN, are tightly related to pain induced negative affect. Moreover, the MCLS seems to be involved in the modulation of negative affective states occurring in pain conditions. (Caputi et al., 2019; Jesús David Lorente et al., 2020; Massaly et al., 2019) Thereby, we measured the mRNA levels of pDYN *and* KOR in the NAc and the Amy by qRT-PCR together with its protein expression in these areas by ELISA and western blot.

On one hand, as observed in Figures 3A to 3D, the presence of inflammatory pain in male rats led to some alterations in the pDYN and KOR mRNA expression in the Amy and NAc at the end point of the behavioural characterisation. However, the protein levels remained unaltered (figure 3E to 3H). Indeed, inflammatory pain in male rats increased pDYN mRNA expression in the Amy (figure 3B, p= 0.05), and interestingly reduced KOR mRNA in the NAc (figure 3C, p= 0.026).

**Figure 3.**
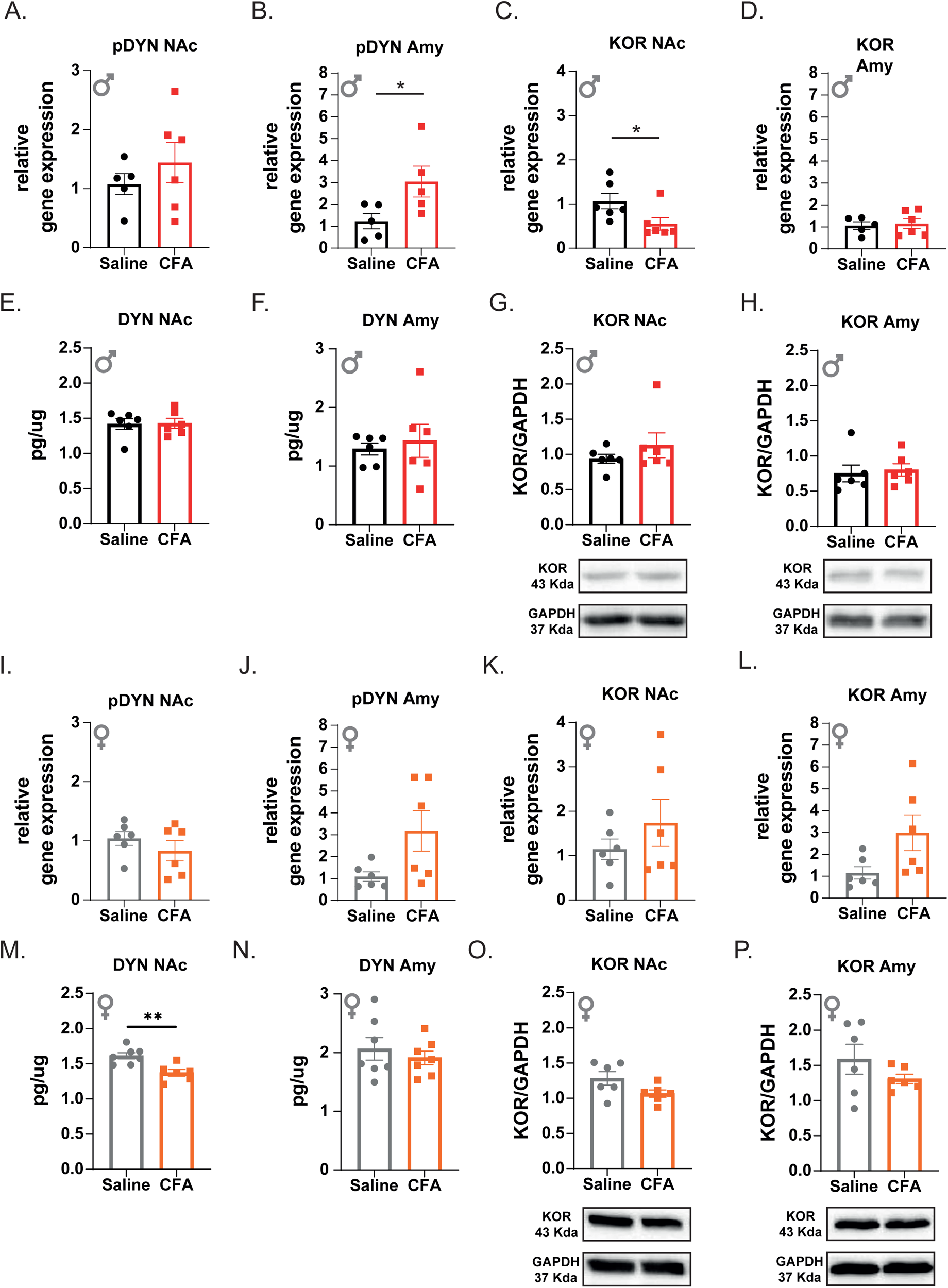
Kappa opioid receptor/Dynorphinergic system alterations induced by inflammatory pain in male and female rats. A) Mean ± SEM of pDYN mRNA relative gene expression in NAc of male rats. B) Mean ± SEM of pDYN mRNA relative gene expression in Amy of male rats. C) Mean ± SEM of KOR mRNA relative gene expression in NAc of male rats. D) Mean ± SEM of KOR mRNA relative gene expression in Amy of male rats. E) Mean ± SEM of DYN protein expression (ug/tissue) in NAc of male rats. F) Mean ± SEM of DYN protein expression (ug/tissue) in Amy of male rats. G) Mean ± SEM of KOR protein expression in NAc of male rats (KOR/GAPDH arbitrary units). H) Mean ± SEM of KOR protein expression in Amy of male rats (KOR/GAPDH arbitrary units). I) Mean ± SEM of pDYN mRNA relative gene expression in NAc of female rats. J) Mean ± SEM of pDYN mRNA relative gene expression in Amy of female rats. K) Mean ± SEM of KOR mRNA relative gene expression in NAc of female rats. L) Mean ± SEM of KOR mRNA relative gene expression in Amy of female rats. M) Mean ± SEM of DYN protein expression (ug/tissue) in NAc of female rats. N) Mean ± SEM of DYN protein expression (ug/tissue) in Amy of female rats. O) Mean ± SEM of KOR protein expression in NAc of female rats (KOR/GAPDH arbitrary units). P) Mean ± SEM of KOR protein expression in Amy of female rats (KOR/GAPDH arbitrary units). * Denotes significant differences between groups (t-student for independent samples or Welch’s t test p < 0.05).

On the other hand, mRNA expression of pDYN and KOR 18 days *after* the inflammatory pain induction, were not significantly altered in the NAc or in the Amy (figure 3I to 3L) of female rats, although some tendencies to increase the mRNA can be perceived. Protein expression also remained unaltered (figure 3N to 3P) regardless of the DYN in the NAc that was significantly reduced in the presence of inflammatory pain (fig 3M, p = 0.003).

### 3.4. Kappa opioid receptor blockade rescues inflammatory pain-induced anxiety-like behaviours in female rats

Previous data have shown that the blockade of the KOR/DYN signalling in the NAc shell was an appropriate strategy to impair the development of negative affect and alcohol-relapse-like behaviour induced by pain (Massaly 2019, Lorente 2021). Thus, we found interesting to pharmacologically analyse the role of NAc KOR in the inflammatory pain-induced negative affective states reported in female rats in the present study (figure 1). The blockade of the KOR locally in the nucleus accumbens shell of female rats (figure 5A and S3A) impacted the behaviour in the LDB without altering the SPT. In fact, in the case of the SPT 48h, the ANOVA only detected differences for the time variable (F(2,130)= 15.259; p= 0.0001; η^2^= 0.19) and the interaction time x pain (F(2,130)= 11.576; p= 0.0001; η^2^= 0.151). Further post-hoc analyses revealed that CFA groups decreased their sucrose consumption compared with saline groups at days 1-2 (p= 0.010), while no differences were found for the other variables and interactions (Figure 4B) including the treatment variable and its interactions (see table 1). The analysis of the SPT of 2h, revealed that the consumption of sucrose increases over the time and the animals whose received NorBNI infusion showed a high sucrose consumption than aCSF groups. Indeed, the ANOVA showed differences for the time and treatment variables (F(2,130)= 51.364; p=0.0001; η^2^= 0.597; and F(1,65)= 5.054; p= 0.028; η^2^= 0.072). However, we did not detect differences for pain variable and the interaction between variables (Figure 4C).

**Figure 4.**
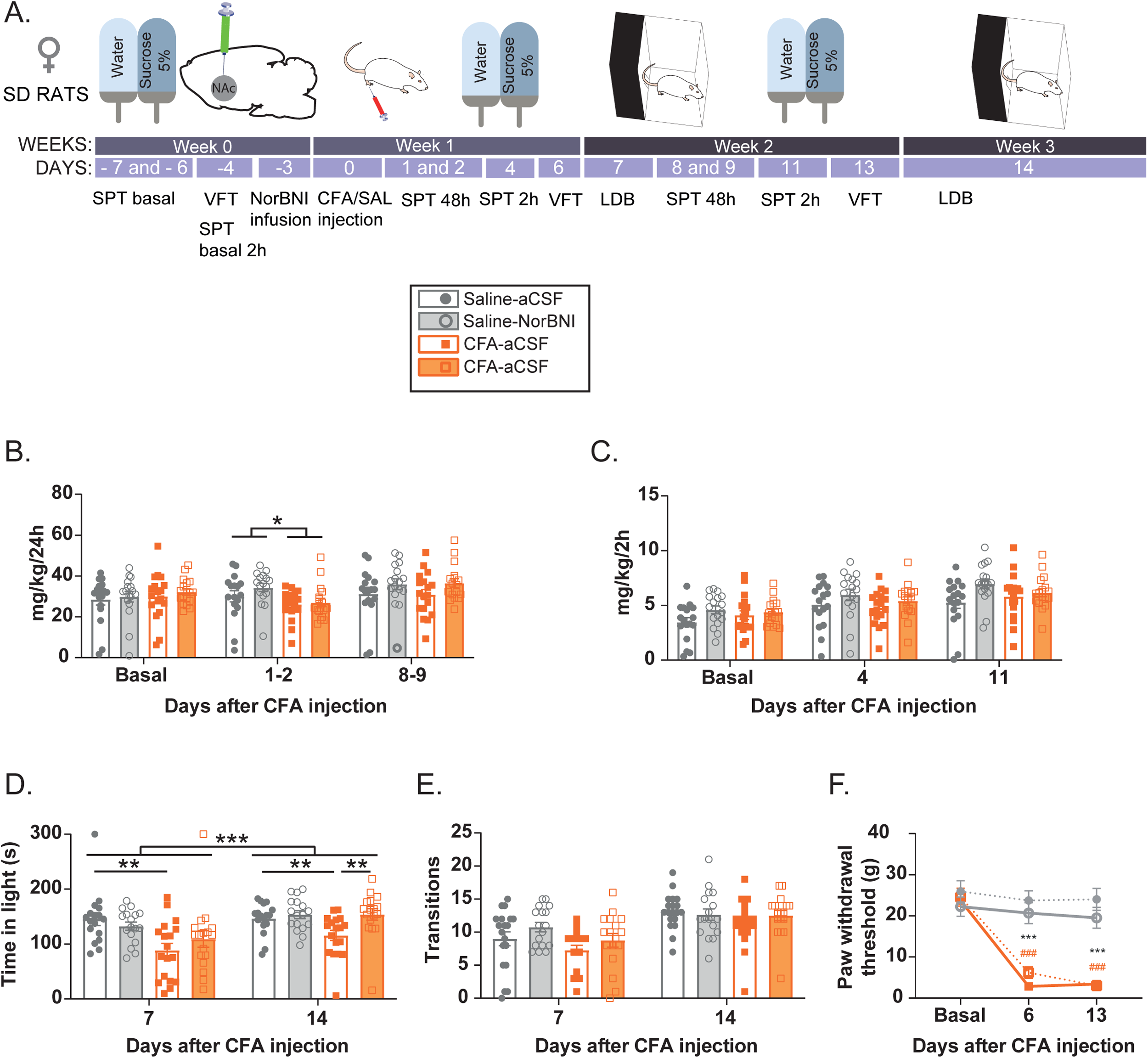
Anhedonia- and anxiety-like behaviour induced by inflammatory pain in female rats is blocked by NorBNI intra-NAc administration. A) Timeline of the experimental design. B) Sucrose consumption during 48 hours in female rats, expressed as mean ± SEM of mg/kg/24 hours mean ± SEM. C) Sucrose consumption during 2 hours in female rats, expressed as mean ± SEM of mg/kg/2 hours. D) Time in light chamber (seconds) in the light-dark box test (LDB). E) Number of transitions between chambers in the LDB, expressed as mean ± SEM. F) Mechanical sensitivity threshold (g) in female rats, expressed as mean ± SEM of paw withdrawal threshol. Grey and orange bars represent saline- and CFA-treated female rats respectively, and empty and full bars represents the intracranial treatments of aCSF and NorBNI respectively. *, ** and *** denotes significant differences between groups (three-way ANOVA for repeated measures followed by Bonferroni multiple comparisons test, p < 0.05, p < 0.01 and P < 0.001, respectively), and #, ## and ### denotes significant differences within groups compared with the basal measures (two-way ANOVA for repeated measures followed by Bonferroni multiple comparisons test, p < 0.05, p < 0.01 and P < 0.001, respectively). Abbreviation: SPT, sucrose preference test; VFT, Von Frey test; CFA, Complete Freund’s Adjuvant; SAL, saline; LDB, light-dark box test; NorBNI, Norbinaltorphimine.

Finally, when analysing the LDB test we observed that NorBNI reverse the decrease in the time spent in light box of CFA female rats without altering the transition between boxes. For the time in light, we detected differences in the time variable (F(1,65)= 12.969, p= 0.001; η^2^= 0.166), showing that animals spent more time in light during the second week. Furthermore, we detected differences in the pain variable (F(1,65)= 11.470, p= 0.001; η^2^= 0.15) and in the interaction between pain x treatment (F(1,65)= 3.954, p= 0.05; η^2^= 0.057). The treatment variable and the other interactions showed no significative differences, and their F values are showed in table 1. Post-hoc analyses of the significant interaction and variables revealed that pain-suffering female rats that received aCSF spent a shorter time in light box, compared to the pain-free animals at 7 and 14 days (p_7_= 0.001, p_14_= 0.012) while the pain-suffering female rats receiving NorBNI spent in the light compartment the same time as saline groups (Figure 4D, p_7_= 0.181; p_14_= 0.974). Thus, the administration of NorBNI counteracted the anxiety-like behaviour induced by CFA injection in female rats. The analysis of the transitions between boxes detected a significant effect for the time (F(1,65)= 45.153; p= 0.0001) but not for the other variables and their interactions (see table1). Thus, the transitions at 7 days were lower than the transitions at 14 days either after saline or after CFA injection.

Finally, we analysed the mechanical sensitivity thresholds with the VFT to confirm that all CFA-treated rats maintained values until the end of the protocol. We detected differences in pain (F(1,65)= 69.774; p= 0.0001; η^2^= 0.518) and time (F(1,65)= 55.861; p= 0.0001; η^2^= 0.521) variables and in the interaction between time x pain (F(1,65)= 36.695); p= 0.0001; η^2^= 0.41), while no differences were detected for treatment variables and for their interactions (see table 1). Post-hoc analyses revealed that female rats showed lower paw withdrawal thresholds after 7 and 14 days following CFA injection compared with their own basal values (before the injection, p_7_= 0.0001); p_14_= 0.0001) and with their saline counterparts (p_7_= 0.0001; p_14_= 0.0001).

## 4. Discussion

Here, we provide new details of the sex- and time-dependent effect of inflammatory pain on the development of negative affective states. Inflammatory pain induces persistant negative affective states in females. Very interestingly, we found biochemical alterations in the HPA axis and in the mRNA of pDYN and KOR in NAc and amygdala in male rats that did not correlate with the behaviour observed. Alterations in protein levels were found only in the case of CFA-female rats, showing a reduction of DYN in NAc, although the pharmacological blockade of the KOR in the NAc effectively blunted the anxiety-like behaviour showed by CFA-female rats. Thus, we propose that alterations in how these systems respond to the presence of inflammatory pain in males *versus* females, might underlie the observed sex-dependent pain-induced negative affect. Although our biochemical analysis is not compelling, the blockade of the KOR within the NAc shell shows a potential target to treat psychological comorbidities of pain conditions.

Many studies highlighted that pain induces anxiety- and anhedonia-like behaviours as well as low motivation for natural reinforcement under a goal-directed behaviour (Lorente et al., 2020; Markovic et al., 2021; Massaly et al., 2019). However, unfortunately, most of these studies used only male subjects and limited observations only to specific time points after pain induction. Therefore, the aim of this study was to fill this gap in the literature by exploring whether inflammatory pain affects anxiety- and anhedonia-like behaviours in both sexes separately to avoid the overseeing of small differences. In addition, we added longer periods of time to study the time course of these pain-induced behaviours. In this framework, the involvement of the HPA axis and KOR/DYN signalling has also been explored in relevant brain areas. Here, we shown that both male and female rats developed anhedonia-like behaviour at the early phase of CFA-induced pain. Indeed, we observed a reduction of sucrose intake on day 1 and 2 after CFA-administration, being the effect more pronounced in female rats. This first finding agrees with previous results indicating a reduction of sucrose consumption a few days after pain induction both in non-operant and operant procedures in rats (Hipolito et al., 2015; Markovic et al., 2021; Massaly et al., 2019; Schwartz et al., 2014). Together with this decrease in the motivation for sucrose consumption, we also found that development of anxiety-like behaviour was affected by pain. However, this specific behaviour was only observed 2 weeks after the induction of inflammatory pain in female rats. Contradictory data regarding pain induced anxiety-like behaviour in rodents are present in literature. A detailed review by Kremer and co-workers, reported that CFA-inflammatory pain induces anxiety-like behaviour, at the onset of the pain condition (1-2 weeks) and after becoming chronic (> 2 weeks); however a high number of studies did not report these anxiety-like behaviour (Kremer et al., 2021). In this regard, it is very important to highlight that most of the examined studies only used male rodents and, the few studies that used female animals showed that CFA was sufficient to induce anxiety-like behaviour (Liu et al., 2019; Pitzer et al., 2019; Refsgaard et al., 2016). In addition, Liu and collaborators also showed sex-dependent differences of pain effects not only at behavioural but also at biochemical level.

It has been well documented that stress and anxiety alter cortisol plasma levels, usually inducing an increase of this hormones. However, in the presence of chronic stress cortisol levels have also been reported to be decreased (Adzic et al., 2009; Costache et al., 2020; Gong et al., 2015; Kim et al., 2018). In the last years, pain has been defined as a stressor that can participate in the development of psychological diseases such as anxiety and depressive disorders (Csupak et al., 2018; Tsang et al., 2008). Accordingly, we analysed cortisol plasma levels after 18 days of pain induction. Interestingly, we did not observe differences in male rats; instead, pain-suffering female rats showed a significant reduction compared with their control counterparts. It is known that plasma cortisol levels increase after acute pain, however a prolonged or exaggerated stress condition in response to pain-and non-pain-related stressors can aggravate pain and promote further disabilities (Edwards et al., 2008; Hall et al., 2011; Heim et al., 2000; Tak & Rosmalen, 2010). Therefore, a desirable response of the stress system in the presence of prolonged pain would be the reduction of its activation towards a homeostatic level, to avoid worsening of the pain condition. This regulation might explain data here observed in male rats that despite of showing some gene expression changes, did not display alterations in the CRF and CRFR1 protein levels in NAc and Amy. In addition, cortisol plasma levels of CFA-male rats were not different from controls 18 days after pain induction.

On the other hand, the chronic reactivation of stress responses leads to exhaustion of HPA axis and induces hypocortisolaemia (Penninx et al., 2007), that may explain the cortisol dysfunction that has been related with idiopathic pain and inflammation responses (Hannibal & Bishop, 2014). Furthermore, hypocortisolism has been also related to chronic stress and pain disorders such as fibromyalgia among others (Ehlert et al., 2001; Tak & Rosmalen, 2010; Tsigos & Chrousos, 2002). Therefore, our data suggest that pain when suffered by female rats produces a higher response of the HPA axis than in males leading to a hypocortisolism condition together with a lack of adaptation of CRF and CRFR1 18 days after the CFA administration, in the NAc and the Amygdala.

In the research framework of pain related behaviours, several studies investigated CRF and KOR/DYN systems and their relationships, in different areas of the MCLS (Hein et al., 2021; Massaly et al., 2019; Mousa et al., 2007; Navratilova et al., 2019; Palmisano et al., 2019). Since these studies reported the ability of KOR antagonism to reduce CRF release and to blunt pain-related behaviors, we decided to further investigate the biochemical adaptations on the KOR/DYN system in the NAc and Amy, in our experimental conditions. Although we are aware of limitation entailed by analysing the gene and protein expression only 18 days after pain induction, our pharmacological data confirm the involvement of KOR/DYN signalling in NAc in pain-induced anxiety-like behaviour. Here reported data showing alterations in the gene and protein expression of DYN and KOR in both males and females. In the case of males, alterations in gene expression of KOR without effect in the total protein have been observed. Accordingly, the downregulation of KOR gene expression has been previously described in the sciatic nerve chronic constriction model (Palmisano et al., 2019). Two explanations are plausible; first, the downregulation of KOR could be related with the dynorphin hyperactivity induced by pain of in this area (Massaly et al., 2019) at early stages when animals showed a tendency to decrease in the sucrose intake. Secondly, it should be underlined that gene expression changes are not invariably related with protein levels (Maier et al., 2009), and this ability to regulate the final expression of the functional protein might explain the lack of pain-induced anxiety-like behaviours in male rats.

Interestingly at 18 days after CFA injection, pain-suffering female rats showed a reduction in dynorphin peptide content in the NAc compared with control counterparts. It has been demonstrated that pain induces an increase in DYN levels in NAc 2 days after the pain induction and that KOR blockade with NorBNI prevents pain-related behaviour (Massaly et al., 2019); accordingly, the reported results here of pharmacological KOR inactivation support this notion. Therefore, DYN reduction in the NAc of CFA-treated female rats could result from of a late neuroadaptation mechanism aiming the recovery of homeostasis. Nonetheless, further experiments detailing the dynamics of the alterations in both the CRFR1/CRF and KOR/DYN signalling and the HPA axis underlying the occurrence of anhedonia and negative affect in the context of pain in male and female animals are warranted.

In conclusion, we describe anhedonia- and anxiety-like behaviours derived from the development of an inflammatory pain condition that depends on the sex and the time course of the pain condition. Indeed, anhedonia-like behaviour is detected in both male and female rats at early phase, whereas anxiety-like behaviour is only observed in female rats when the chronic inflammatory pain is stablished. Dynamic alteration of KOR/DYN and CRFR1/CRF systems, along with cortisol plasma levels, might underlie these observed behavioural adaptations, nonetheless further detailed studies are warranted. Finally, the pharmacological blockade of KOR in the NAc shell of CFA-treated female rats impairs the development of anxiety-like behaviours in these animals, supporting the role of this system in pain-induced negative affective state in female rats.

## Acknowledgements

This study has been supported by Spanish Ministerio de Ciencia e Innovación PID2019-109823RB-I00 (LJH), Spanish Ministerio de Sanidad, Delegación del Gobierno para el Plan Nacional sobre Drogas PNSD2019I038 (LH), Italian grants PRIN2017SXEXT5_005 to PR; Grant RFO2020 to SC. Javier Cuitavi is supported by a Atracció de Talent PhD Fellowship from the University of Valencia (UV-INV-PREDOC-1327981).

We would like to thank Ms. Pilar Laso, Ms. Luisa Goicoechea and Staff from the “Servei d’Investigació-UV” for grant management. We would also thank Dr. Inmaculada Noguera, Chief Veterinarian Officer, and the personnel of the Animal facilities (SCSIE) at the University of Valencia for their help and effort in assuring animal welfare.

## 6. Figure legends

**Table 1. P values of the graphs presented.**

**Figure S1. Daily calendar of the behavioural protocol.** CFA, complete Freund adjuvant; SAL, saline; LDB, light–dark box test; SPT, sucrose preference test; VFT, von Frey test.

**Figure S2. Schematic of the brain areas dissection.** A) Area of the NAc dissection B) Area of the amygdala dissection.

**Figure S3. Daily calendar of the behavioural protocol of the Experiment 2 and intracranial administration verification.** A) Daily calendar of the behavioural protocol. B) Dots represent the site where aCSF and NorBNI were injected: Black represent saline/aCSF treated female rats, grey represent saline/NorBNI treated female rats, red represents CFA/aCSF treated female rats, and orange represents CFA/NorBNI treated female rats. Abbreviatures: CFA, complete Freund adjuvant; SAL, saline; LDB, light–dark box test; SPT, sucrose preference test; VFT, von Frey test.

