## Supplementary material for "Sex-dependent effect of inflammatory pain on negative affective states is prevented by kappa opioid receptors blockade in the nucleus accumbens shell": Table 1

| Figures | | Variables | F | p value |
|---|---|---|---|---|
| 1 | B | Pain | F (1, 22) = 0.069 | p = 0.795 |
| | | Time | F (3, 66) = 6.636 | p = 0.001 |
| | | Time*Pain | F (3, 66) = 3.709 | p = 0.016 |
| | C | Pain | F (1, 25) = 7.688 | p = 0.01 |
| | | Time | F (3, 75) = 3.496 | p = 0.02 |
| | | Time*Pain | F (3, 75) = 7.819 | p < 0.001 |
| | D | Pain | F (1, 22) = 0.302 | p = 0.558 |
| | | Time | F (3, 66) = 8.886 | p < 0.001 |
| | | Time*Pain | F (3, 66) = 0.597 | p = 0.448 |
| | E | Pain | F (1, 25) = 2.034 | p = 0.166 |
| | | Time | F (3, 75) = 1.593 | p = 0.198 |
| | | Time*Pain | F (3, 75) = 0.593 | p = 0.621 |
| | F | Pain | F (1, 22) = 0.108 | p = 0.746 |
| | | Time | F (1, 22) = 19.284 | p < 0.001 |
| | | Time*Pain | F (1, 22) = 0.926 | p = 0.346 |
| | G | Pain | F (1, 25) = 7.287 | p = 0.012 |
| | | Time | F (1, 25) = 4.355 | p = 0.047 |
| | | Time*Pain | F (1, 25) = 0.031 | p = 0.862 |
| | H | Pain | F (1, 22) = 0.289 | p = 0.596 |
| | | Time | F (1, 22) = 18.053 | p < 0.001 |
| | | Time*Pain | F (1, 22) = 0.03 | p = 0.863 |
| | I | Pain | F (1, 25) = 0.031 | p = 0.862 |
| | | Time | F (1, 25) = 13.782 | p = 0.001 |
| | | Time*Pain | F (1, 25) = 0.04 | p = 0.843 |
| | J | Pain | F (1, 22) = 6.223 | p = 0.021 |
| | | Time | F (2, 44) = 4.148 | p = 0.022 |
| | | Time*Pain | F (2, 44) = 11.538 | p < 0.001 |
| | K | Pain | F (1, 25) = 56.140 | p < 0.001 |
| | | Time | F (2, 50) = 3.252 | p = 0.047 |
| | | Time*Pain | F (2, 50) = 11.783 | p < 0.001 |
| 2 | A | Pain | | p = 0.045 |
| | B | Pain | | p = 0.038 |
| | C | Pain | | p = 0.075 |
| | D | Pain | | p = 0.046 |
| | E | Pain | | p = 0.999 |
| | F | Pain | | p = 0.093 |
| | G | Pain | | p = 0.589 |
| | H | Pain | | p = 0.445 |
| | I | Pain | | p = 0.177 |
| | J | Pain | | p = 0.394 |
| | K | Pain | | p = 0.42 |
| | L | Pain | | p = 0.818 |
| | M | Pain | | p = 0.590 |
| | N | Pain | | p = 0.033 |
| 3 | A | Pain | | p = 0.392 |
| | B | Pain | | p = 0.05 |
| | C | Pain | | p = 0.026 |
| | D | Pain | | p = 0.755 |
| | E | Pain | | p = 0.926 |
| | F | Pain | | p = 0.648 |
| | G | Pain | | p = 1 |
| | H | Pain | | p = 0.485 |

| | | | | |
|---|---|---|---|---|
| | I | Pain | | p = 0.337 |
| | J | Pain | | p = 0.065 |
| | K | Pain | | p = 0.328 |
| | L | Pain | | p = 0.077 |
| | M | Pain | | p = 0.003 |
| | N | Pain | | p = 0.505 |
| | O | Pain | | p = 0.085 |
| | P | Pain | | p = 0.259 |
| | | Pain | F (1, 65) = 0.303 | p = 0.584 |
| | | Time | F (2, 130) = 15.259 | p < 0.001 |
| | | Treatment | F (1, 65) = 2.011 | p = 0.161 |
| | | Time*Pain | F (2, 130) = 11.576 | p < 0.001 |
| | | Time*Treatment | F (2, 130) = 2.203 | p = 0.115 |
| | | Pain*Treatment | F (1, 65) = 0.018 | p = 0.895 |
| | B | Time*Pain*Treatment | F (2, 130) = 1.168 | p = 0.314 |
| | | Pain | F (1, 65) = 0.091 | p = 0.764 |
| | | Time | F (2, 130) = 51.364 | p < 0.001 |
| | | Treatment | F (1, 65) = 5.054 | p = 0.028 |
| | | Time*Pain | F (2, 130) = 1.465 | p = 0.235 |
| | | Time*Treatment | F (2, 130) = 0.397 | p =0.673 |
| | | Pain*treatment | F (1, 65) = 1.238 | p = 0.27 |
| | C | Time*Pain*Treatment | F (2, 130) = 0.987 | p =0.375 |
| | | Pain | F (1, 65) = 11.470 | p = 0.001 |
| | | Time | F (1, 65) = 12.969 | p = 0.001 |
| | | Treatment | F (1, 65) = 2.590 | p = 0.112 |
| | | Time*Pain | F (1, 65) = 3.583 | p = 0.063 |
| | | Time*Treatment | F (1, 65) = 2.043 | p = 0.158 |
| | | Pain*Treatment | F (1, 65) = 3.954 | p = 0.05 |
| | D | Time*Pain*Treatment | F (1, 65) = 0.006 | p = 0.937 |
| | | Pain | F (1, 65) = 4.451 | p = 0.039 |
| | | Time | F (1, 65) = 45.153 | p < 0.001 |
| | | Treatment | F (1, 65) = 2.150 | p = 0.147 |
| | | Time*Pain | F (1, 65) = 0.745 | p = 0.391 |
| | | Time*Treatment | F (1, 65) = 1.548 | p = 0.218 |
| | | Pain*treatment | F (1, 65) = 4.522 | p = 0.596 |
| | E | Time*Pain*Treatment | F (1, 65) = 0.926 | p = 0.339 |
| | | Pain | F (1, 65) = 69.774 | p < 0.001 |
| | | Time | F (1, 65) = 55.861 | p < 0.001 |
| | | Treatment | F (1, 65) = 0.751 | p = 0.389 |
| | | Time*Pain | F (1, 65) = 36.695 | p < 0.001 |
| | | Time*Treatment | F (1, 65) = 0.565 | p = 0.57 |
| | | Pain*Treatment | F (1, 65) = 3.029 | p = 0.087 |
| 4 | F | Time*Pain*Treatment | F (1, 65) = 0.131 | p = 0.877 |
