## Supplementary figures and images for "Sex-dependent effect of inflammatory pain on negative affective states is prevented by kappa opioid receptors blockade in the nucleus accumbens shell"

### fig S2

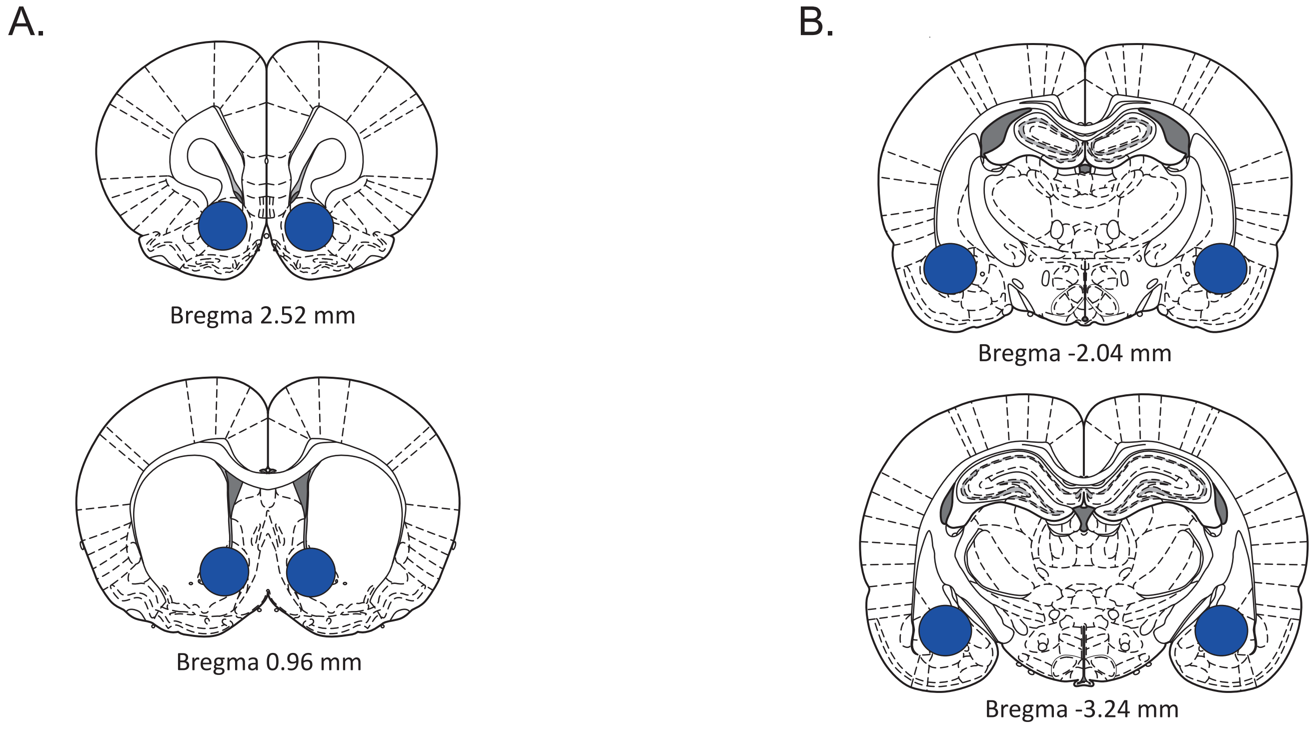
