## Supplementary material for "Sex-dependent effect of inflammatory pain on negative affective states is prevented by kappa opioid receptors blockade in the nucleus accumbens shell": fig S3

A.

| | DAYS | | | | | | |
|---|---|---|---|---|---|---|---|
| WEEK 0 | -7 SPT24h | -6 SPT24h | -5 | -4 SPT2h VFT | -3 Surgery (NorBNI infusion) | -2 | -1 |
| WEEK 1 | 0 CFA/SAL injections | 1 SPT24h | 2 SPT24h | 3 | 4 SPT2h | 5 | 6 VFT |
| WEEK 2 | 7 LDB | 8 SPT24h | 9 SPT24h | 10 | 11 SPT2h | 12 | 13 VFT |
| WEEK 3 | 14 LDB Sacrifice | | | | | | |

B.

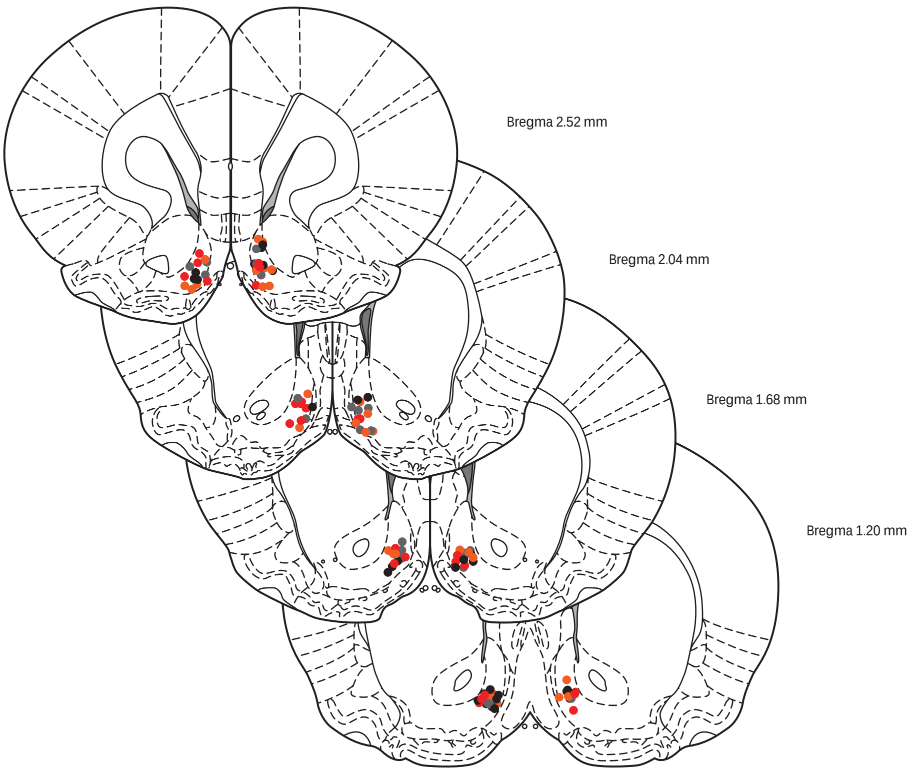
